# Efficient transcriptome profiling across the malaria parasite erythrocytic cycle by flow sorting

**DOI:** 10.1101/2020.11.10.377168

**Authors:** Aliou Dia, Catherine Jett, Marina McDew-White, Xue Li, Timothy J.C. Anderson, Ian H. Cheeseman

## Abstract

*Plasmodium falciparum* is the most virulent and widespread of the human malaria parasite species. This parasite has a complex life cycle that involves sexual replication in a mosquito vector and asexual replication in a human host. During the 48-hour intraerythrocytic developmental cycle (IDC), parasites develop and multiply through the morphologically distinct ring, trophozoite and schizont stages. Stage-specific transcriptomic approaches have shown gene expression profiles continually change throughout the IDC. Cultures of tightly synchronized parasites are required to capture the transcriptome specific to a developmental stage. However, the most commonly used synchronization methods require lysis of late stages, potentially perturbing transcription, and often do not result in tightly synchronized cultures. To produce complete transcriptome profiles of the IDC a synchronous culture requires frequent sampling over a 48-hour period, this is both time consuming and labor intensive. Here we develop a method to sample the IDC densely by isolating parasites from an asynchronous culture with fluorescence activated cell sorting (FACS). We sort parasites in tight windows of IDC progression based on their DNA/RNA abundance. We confirmed the tight synchronization and stage specificity by light microscopy and RNAseq profiling. We optimized our protocol for low numbers of sorted cells allowing us to rapidly capture transcriptome profiles across the entire IDC from a single culture flask. This methodology will allow malaria stage-specific studies to perform experiments directly from asynchronous cultures with high accuracy and without the need for labor-intensive time-course experiments.

## Introduction

Since the publication of the malaria parasite genome [1], transcriptome analysis has become an important tool in malaria research. Many stage specific analyses of transcriptomes from different malaria parasite species have shown a highly regulated “just in time” gene expression program accompanying the transition between stages and revealed discrete transcriptional programs within the intraerythrocytic developmental cycle (IDC) [2–8]. While study of the IDC transcriptome has led to major progress in understanding malaria biology, investigation of transcript levels between different stages of parasites remains cumbersome due to the asynchronous nature of *Plasmodium* parasites in culture and the major efforts required for synchronizing and collecting samples during the 48-hour duration of the IDC. In order to capture the transcriptome specific to a particular developmental stage it is essential to use cultures of tightly synchronized parasites

. Various methods of synchronization of *P. falciparum* have been developed which rely on either selective killing of parasites (i.e. by sorbitol treatment [9]), isolation of trophozoite/schizont stages (i.e. by magnetic column separation [10] or concentration by Percoll gradient [11]) or reinforcing of natural lifecycle rhythms by temperature shifts [12]. No method provides absolute synchrony. For instance, in an asynchronous culture sorbitol treatment kills only the trophozoite stage, requiring two rounds of sorbitol treatment for 90% synchronicity [13]. Percoll density gradients can increase stage specific purity of the synchronized culture, though also contain sorbitol [14, 15] which has a toxic effect on the parasite and can cause stress on living cells. Since *Plasmodium* species have stage specific transcriptional programs, imperfectly synchronized samples can lead to confounding interpretations from analysis of mixed stages instead of a discrete single stage [16]. Even in the case of a well synchronized culture, an additional challenge for standard approaches is the need to frequently sampling over a 48-hour period to produce the complete transcriptome profile of the IDC. Such approaches typically require nanogram quantities for RNAseq library preparation further limiting the scalability of standard approaches.

To alleviate the need for synchronization and eliminate the requirement for 48 hours of sampling, we developed a flow cytometric sampling protocol. Flow cytometry techniques have previously been used to identify *Plasmodium* parasite developmental stages by staining both the DNA and the RNA of the parasites [17, 18]. Here, we developed an improved method we call pFACS-RNAseq (*Plasmodium*-FACS-RNAseq) to capture the whole life cycle of malaria parasites in tight windows with high stage specificity. The identified stages are subdivided in multiple gates or populations for higher temporal resolution of RNAseq profiling throughout the 48-hour IDC. Using our pFACS-RNAseq approach we decreased the sampling time of the time-course from 6 days (including multiple synchronization steps and 2 days of sampling) to 2 hours using fluorescence-activated cell sorting (FACS). RNA extraction is not required for our method, and cDNA synthesis occurs in the same 96 well plate into which the cells were sorted, saving an additional 2-4 hours. After cDNA synthesis, the samples are bead cleaned and amplified to produce sufficient cDNA for library preparation in optimized reaction volumes. Our final cost for this workflow is $15 per sample (sorted well), compared to ~$45 for traditional RNA isolation, cDNA synthesis, and library preparation. Notably, this does not take into account the associated culture costs, or costs associated with hands on time in the traditional protocol. Our protocol will enable transcriptome profiling of cultivable species and also promises to be highly beneficial for species that are only amenable to brief *ex vivo* culture, such as *P. vivax.*

## Results

### Identification of malaria parasites asexual stages by FACS

The genus *Plasmodium* doesn’t follow the conventionalG1, S, G2 and M phases of the standard eukaryotic cell cycles [19]. Progression through the intraerythrocytic development cycle (IDC) occurs over 48 hours, and is accompanied by progressive, asynchronous genome replication [20]. The IDC is divided into three major morphologically distinct stages (the ring, trophozoite and schizont), with bell shaped transcriptional activity starting at the early ring stage then increases at late ring before peaking in the early and late trophozoite stages. Finally, the transcriptional activity decreases again when the parasite mature into schizont stages [21]. The canonical morphological stages of the IDC can be detected by flow cytometry using DNA and RNA stains in both fixed and unfixed cells [17, 18]. We have previously used live cell DNA dyes to distinguish between uninfected and infected erythrocytes of different stages for single genome sequencing [22]. Change in DNA content and transcriptional activity are not perfectly correlated. Sims *et al* demonstrated the transcriptional activity for a given stage does not increase linearly with the average number of nuclei in parasites of that stage [23]. We hypothesized the addition of an RNA-specific dye would improve the resolution to which life cycle stages can be isolated. To test this, we used live cell dyes for DNA (VybrantDye Violet™) and RNA (SYTO RNA Select™) to stain an asynchronous culture of *P. falciparum* (Fig. 1A). As parasites are cultivated in anucleated human red blood cells (RBCs), there is little host nucleic acid to interfere with this assay. A dot plot of VybrantDye Violet™versus SYTO RNA Select™replicates the imperfect correlation between DNA and RNA content (Fig. 1B). As expected, fluorescence in the DNA channel demonstrated clear separation between uninfected and infected cells. We observe a clear trajectory of change in nucleic acid composition which fits the known developmental progression of the IDC (Fig. 1B). First, transcriptional activity increases without an increase in genome copy number as parasites develop from ring to trophozoite stages after RBC invasion. This is captured by increased SYTO fluorescence (but not VybrantDye fluorescence). Next DNA replication proceeds, along with a moderate increase in RNA content as parasites develop into schizont stages, this is captured by increases in both VybrantDye and SYTO fluorescence. Based upon this model we identified cell populations putatively corresponding to uninfected erythrocytes, ring, trophozoite and schizont stages (Fig. 1B). To validate our inference, parasites from each putative developmental stage were sorted, Wright-Giesma stained and observed by microscopy (Fig. 1C-E). This confirmed that the sorted gates correspond to the morphologically distinct ring (Fig. 1C), trophozoite (Fig. 1D) and schizont stages (Fig. 1E).

**Fig. 1.**
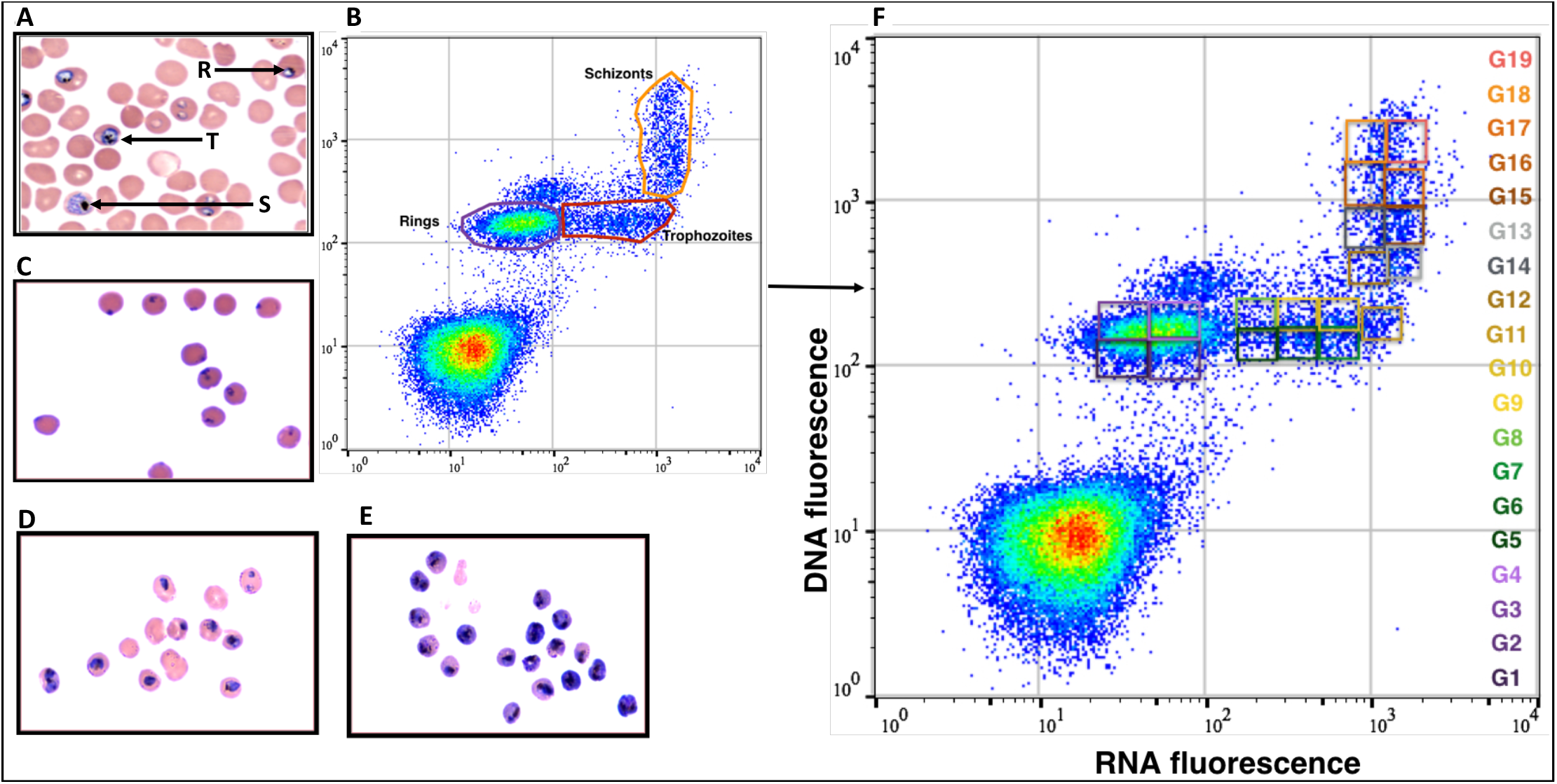
Flow cytometry analysis, gating strategy and cell sorting from an asynchronous *P. falciparum* culture. (A) Wright-Giesma stained thin blood smear of the unsynchronized culture of P. falciparum used for flow sorting. In this representative field, ring, trophozoite and schizont stages are visible (marked R, T and S). (B) Flow cytometry dot plots of the unsynchronized culture stained by DNA and RNA live cell dyes. Uninfected erythrocytes are clearly separated from infected erythrocytes by their DNA and RNA content. We identified 3 major populations of cells. Wright-Giesma stained smears from gates of the 3 cell populations revealed morphologies corresponding to ring (C), trophozoite (D) and schizont (E) stages. (F) The 3 main stages of P. falciparum profiled by flow cytometric were partitioned in 19 gates (G1 to G19) for initial RNA-seq profiling.

### Dense sampling of the parasite IDC

Our primary goal is to develop a FACS approach to allow high throughput transcriptomics of the parasite IDC. To test the resolution we achieve across the IDC, we divided the 3 morphological stages detected by the flow cytometry into 25 gates denoted P1 to P25. For each gate 100 cells were sorted in triplicate into 96 well plates for RNAseq profiling using the QIAGEN FX Single Cell RNA Library Kit (QIAGEN). This kit was developed for RNAseq from single cells or low amounts of RNA and uses a PCR-free whole transcriptome amplification protocol to reduce bias from the PCR. We sequenced each of these 75 libraries to high coverage, generating a mean of ~1.7 million reads (range 0.9-3.2M reads) per library. After aligning reads to the *P. falciparum* genome, we saw a very low proportion of uniquely mapped reads (mean of 17.85% SD: 0.0911) (Fig. 2A and Supplementary Table 1)) and a considerable number of unmapped reads (mean of 51% (SD: 0.200) (Fig. 2B and Supplementary Table 1)). Using a threshold of TPM>1 reads per gene we detected 3,904 genes per gate, with gates from ring stages (P1-P8) showing a mean of 4,062 genes (range 2,665-5,073), gates from trophozoite stages (P9-P16) showing a mean of 3,573 genes (range (3,128-4,903) and gates from schizont stages (P17-P25) showing a mean of 4063 genes (range 2,229-5,479). We detected moderate agreement between replicate samples (Fig. S1 (mean *r^2^* 0.852, range 0.001 - 0.999)) suggesting that despite identifying high numbers of genes, we do not reliably quantify the expression level of these genes. Using multidimensional scaling (MDS) we were able replicate progression through the cell cycle (Fig. 3A) and we correlated the gene expression level of each gate to a ‘gold standard’ dataset generated from synchronized parasites and sampled for 56-hours with 4 hour intervals [24]. This supported the IDC progression of gates P1-P25 through the cell cycle (Fig. 3A (QIAGEN)).

**Fig. 2.**
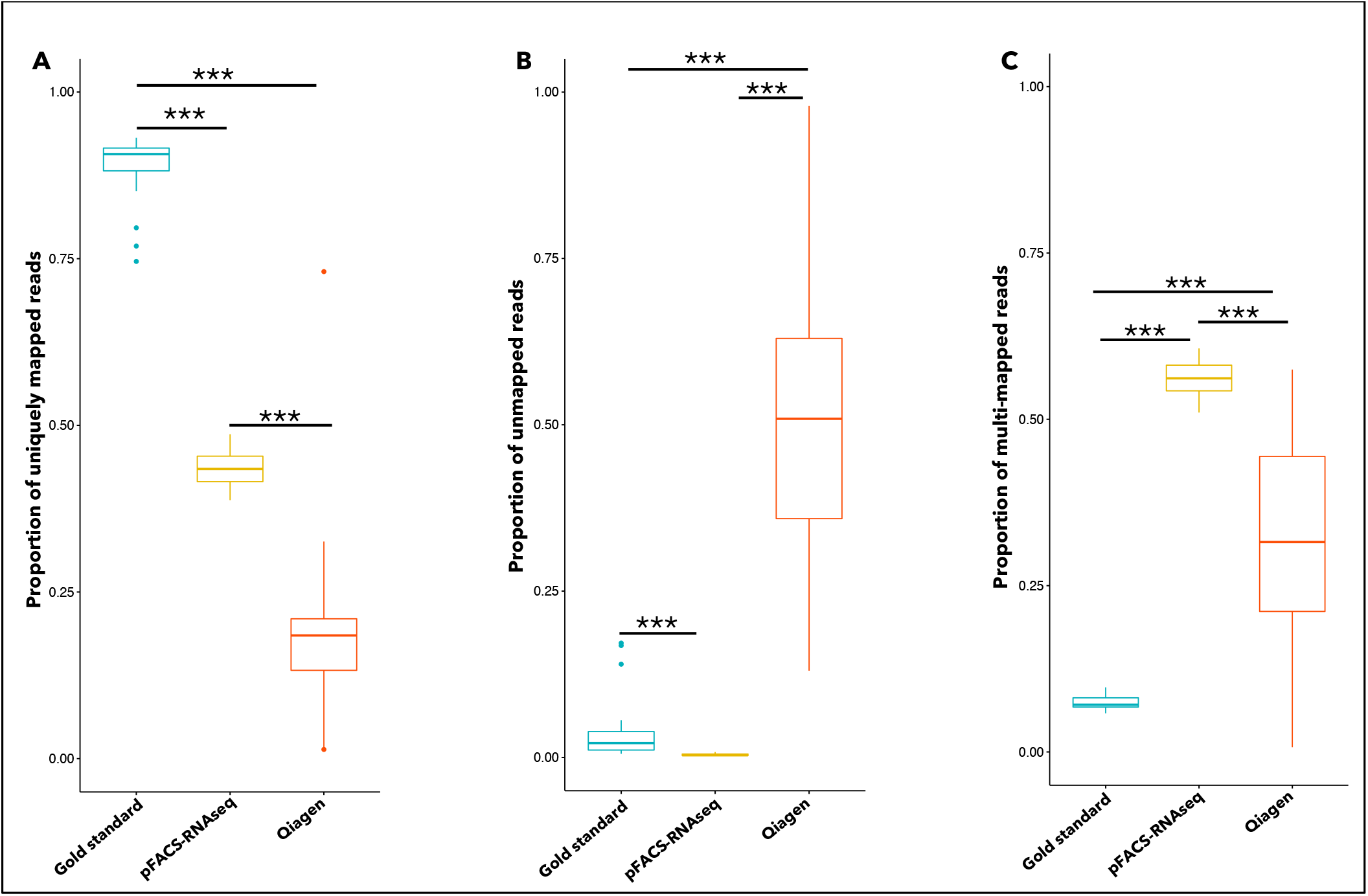
Summary of mapping statistics for each protocol, each proportion here is estimated from all reads detected by the aligner STAR. (A) Average proportion of uniquely mapped reads to the 3D7 Plasmodium falciparum reference genome. (B) Proportion of unmapped reads: these reads did not align to the 3D7 P. falciparum reference genome. (C) Proportion or reads that mapped equally well at more than one locus (multi-mapped reads). Kruskal-Wallis global comparison shows significant differences between the approaches. Significance levels from Pairwise comparisons using Wilcoxon rank sum test with continuity correction are represented on the Figure (***p≤0.001).

**Fig. 3.**
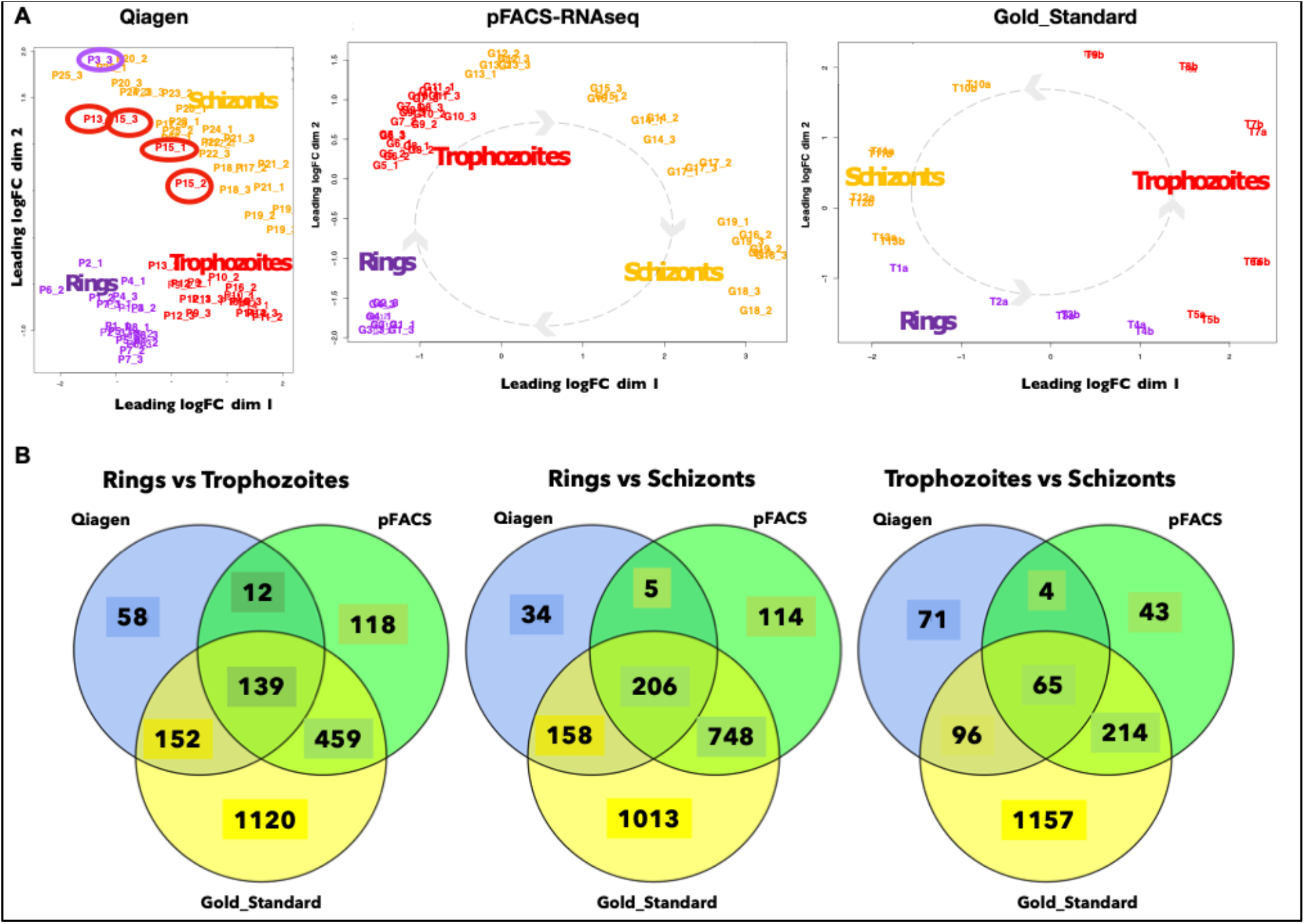
Concordance of gene expression analysis throughout *P. falciparum* life cycle. (A) Multidimensional scaling (MDS) performed from the QIAGEN, pFACS-RNAseq and the Gold standard protocols. (B) Venn diagram showing the number of differentially expressed genes shared between the three protocols (pFACS: pFACS-RNAseq).

### Optimization of transcriptome profiling

Our initial interrogation of the IDC using FACS coupled with RNAseq demonstrated the ability to extract stage specific transcriptomes. However, the cost, coverage and quality of the data need further optimization for routine use. Previous studies [8] have shown the value in optimizing single cell transcriptomics for the highly AT-rich genome of *P. falciparum.* We sought to improve the low input requirements of FACS coupled RNAseq protocol using the molecular crowding single-cell RNA barcoding and sequencing (mcSCRB-Seq) protocol [25]. This approach significantly improves cDNA yield by addition of polyethylene glycol (PEG 8000). Our initial attempts to implement this protocol did not yield successful RNAseq libraries, likely due to the extreme nucleotide bias of the *P. falciparum* genome [1], the lower number of transcripts per cell than human cells and extreme difference in transcript abundance across the cell cycle. We therefore optimized the sequence of the primers by fusing the barcode primer from Quartz-seq [26] to a shortened oligo-dT. We then tested the impact of PEG 8000 on lysis and amplification reactions by measuring PCR efficiency using qPCR. We sorted 3000 ring stage parasites into single wells of a 96-well plate and performed transcriptome amplification either in the presence or absence of PEG 8000. We reached a plateau of the amplification curve after 23 cycles of PCR in the PEG positive wells, and after 28 cycles in the PEG negative wells (Table S5).

In order to streamline processing of reactions for balanced library preparation we identified the numbers of cells from each gate which would result in a similar yield of amplified library. A range of inputs between 1 and 3,000 cells were tested. We used C_T_ values from qPCR to determined that 3000 rings, 200 trophozoites, and 200 schizonts yielded an approximately equivalent cDNA yield (Fig. S4A-C) and implemented this strategy to allow sample pooling without measuring the cDNA content before amplification, as the content was below detectable limits. Based upon these optimizations we identified an optimized protocol which we name pFACS-RNAseq for *Plasmodium-FACS-RNAseq*.

We selected 19 of the previous 25 gates, denoted G1 to G19, as the most informative for the parasite IDC. For each gate, 3000 cells were sorted for the ring stage and 200 cells for both trophozoites and schizonts in triplicate into 96 well plates. cDNA synthesis with PEG and oligo-dT’s were performed in the same sort plate. Additionally, we chose to perform full-length cDNA library preparation with KAPA HyperPlus instead of tagmentation to maximize transcript information.

### Comparison of pFACS-RNAseq

We compared pFACS-RNAseq to RNAseq from both QIAGEN and the gold standard method based on the mapping summary statistics (Table 1) that is an overall indicator of RNAseq data quality. To evaluate the capability of each protocol here, we estimated the percentage of reads that mapped to the reference genome and the unmapped reads for each method. For the gold standard data, more than 96% or the sequenced reads mapped to *P. falciparum* reference genome with more than 88% of uniquely mapped reads and 7% of reads that mapped to multiple location into the reference genome (Table 1).

**Table 1.** Mapping statistics printed from the aligner STAR. For each method used in this paper we have the number of analyzed samples (count), the average of uniquely mapping reads, multi-mapping reads and reads that did not map to the 3D7 reference genome.

| Methods | Count (n) | Mean of Uniquely mapping reads (Sd) | Mean of unmapped reads (Sd) | Mean of multi-mapped reads (Sd) |
| --- | --- | --- | --- | --- |
| **Gold standard** | 26 | 0.8873 (0.0477) | 0.0380 (0.0472) | 0.0747 (0.0111) |
| **pFACS-RNAseq** | 56 | 0.4345 (0.0239) | 0.0040 (0.00153) | 0.5615 (0.0236) |
| **QIAGEN** | 74 | 0.1785 (0.0911) | 0.5133 (0.200) | 0.3082 (0.145) |

The pFACS-RNAseq approach outperformed the QIAGEN approach with 43.45% (SD: 0.0239) of reads mapping to a unique location into the genome (Fig. 2A, Table 1), while 0.40% (SD: 0.00153) of the reads did not map to the *P. falciparum* reference genome (unmapped reads)(Fig. 2B, Table 1) and the remainder of the reads (56% (SD: 0.0236)) mapped to multiple loci on the reference genome (Fig. 2C, Table 1). For the all the mapping statistics indexes, the differences between the 3 methods are significantly different (for the uniquely mapped reads (Kruskal-Wallis chi-squared = 127.05, df = 2, p-value < 2.2e-16); multi-mapped reads (Kruskal-Wallis chi-squared = 120.02, df = 2, p-value < 2.2e-16); unmapped (Kruskal-Wallis chi-squared = 129.55, df = 2, p-value < 2.2e-16). Also, all the pairwise Wilcox test for multiple comparison between methods has shown significant pairwise differences between methods (Tables S1; S2 and S3). Overall, we did not see a substantial improvement in the number of genes detected compared to the QIAGEN protocol which detected in average 3904 genes (range: 2229 - 5479) versus the pFACS-RNAseq that detected 2760 genes in average (range: 1537 - 4398), however, the improvements in reproducibility and cost are considerable.

Previous studies in different *Plasmodium* species revealed that gene expression is regulated in a cascade, with genes differentially expressed as the parasites develop throughout the 48-hour life cycle. To explore the results from the three protocols used in this paper, we performed multidimensional scaling (MDS) to reveal the overall structure of the data. This unsupervised clustering analysis (Fig. 3A) shows that in the MDS generated from the QIAGEN protocol, the technical replicates don’t uniformly cluster together; subsequently it is difficult to effectively produce life cycle transcriptomics with data generated from this protocol as it suggests erroneous mixtures between stages. The MDS made from data generated by the pFACS-RNAseq recapitulated the entire IDC with same pattern as the gold standard data. Indeed, both the gold standard and the pFACS-RNAseq revealed there are 3 mains groups corresponding respectively to rings, trophozoites and schizonts. Additionally, for both these methods, the technical replicates cluster together (Fig. 3A), highlighting the reproducibility and accuracy of pFACS-RNAseq. These results suggest that pFACS-RNAseq is able to capture sufficient transcriptional activities to differentiate not only between stages but also within *P. falciparum* asexual stages.

Given the vastly different library preparation methods we do not expect a direct correlation between gene expression levels between methods to be informative. However, we would expect that each method should internally capture the well described patterns in changing transcript abundance across the IDC. To address this, we identified differentially expressed genes (DEG) to explore gene expression patterns throughout the life cycle. We identified time points well represented in all methods which broadly correspond to ring, trophozoite and schizont stages. We performed pairwise comparisons between each life cycle stage, identifying DEGs significant at p<0.05 after correction for false discovery. In Fig. 3B, the Venn diagrams show the DEGs shared between the three protocols in this study. The pFACS-RNAseq share 3.5 times more DEGs with the gold standard data than the QIAGEN dataset for all the transitions with 1,421 shared DEGs between pFACS-RNAseq and gold standard data, and 406 shared DEGs between QIAGEN and gold standard data. The number of shared DEGs between pFACS-RNAseq and gold standard versus QIAGEN and gold standard correspond to 459 versus 152 (rings to trophozoites); 748 versus 158 (rings to schizonts) and 214 versus 96 (trophozoites to schizonts) respectively.

We then explored whether pFACS-RNAseq captures variation across the life cycle at a higher resolution. In order to identify similarities between the gold standard data and the pFACS-RNAseq data across the intra-erythrocytic life cycle, we performed weighted gene coexpression network analysis (WGCNA). For the WGCNA analysis, we used gene expression data from 13 IDC timepoints. All 5360 genes detected by the standard RNAseq method were used for gene co-expression network construction. There were 8 genes clusters detected noted C1 to C8 (Fig. 4A) containing successively 333; 115; 476; 663; 773; 281; 751 and 329 genes what is a total of 3721 genes that fall into the 8 co-expression clusters. Among which, clusters C1 to C3 are highly expressed in ring stage parasites, clusters C4 to C6 are mainly detected in trophozoites; while C7 and C8 are detected in schizonts. Both methods, the gold standard and the pFACS-RNAseq were able to recover the gene expression patterns from all co-expression clusters. Then we compared gene expression patterns between gold standard data and pFACS-RNAseq. Pairwise cluster comparison between the gold standard data and the pFACS-RNASeq demonstrated strong similarities for the timing of gene expression observed (i.e. the genes that are highly expressed at early, middle or late stage for the gold standard data are the same for the pFACS-RNASeq).

**Fig. 4.**
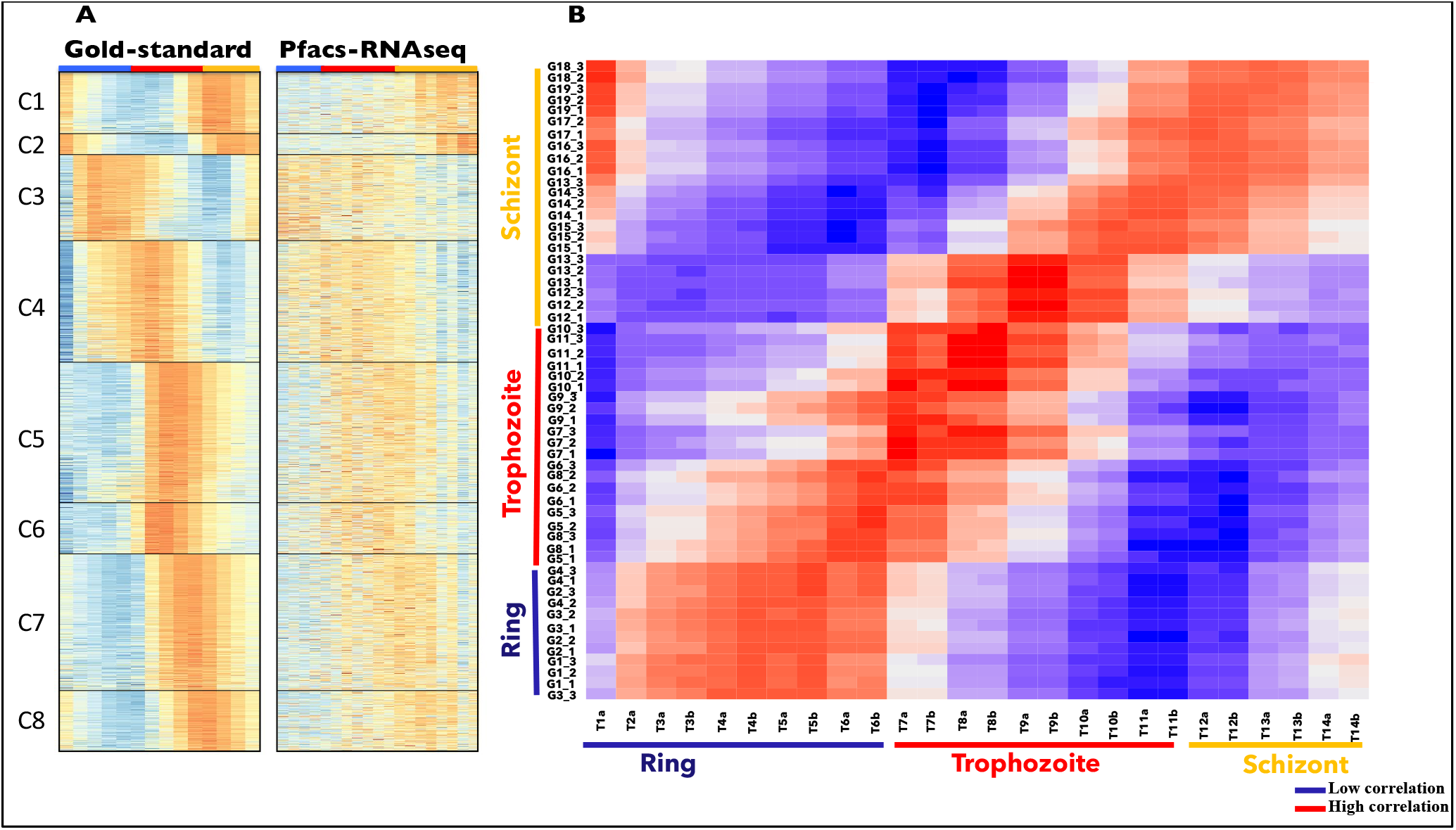
Gene expression similarity analysis based on RNA. (A) Weighted gene coexpression analysis (WGCN) based on gene expression data from 13 timepoints of the gold standard data. All the 5360 genes detected with this method were used for gene co-expression network; 8 clusters were detected (C1-C8). On the heatmaps represented from gene expression levels, brighter orange color indicates higher gene expression while darker blue indicates lower gene expression level. The gold standard and the pFACS-RNAseq methods were both able to reconstruct the gene expression pattern from all co-expression clusters. (B) Heatmap of Pearson correlations between the gold standard gene expression (X-axis) and the pFACS-RNASeq gene expression (Y-axis). Dark blue indicates low correlation while the_dark red indicate high correlation. This heatmap show that both methods are in accordance in terms of gene expression patterns throughout the IDC.

To further explore the accuracy of the pFACS-RNASeq method on the IDC, we calculated Spearman correlation coefficients between gene expression profiles generated by pFACS-RNAseq versus the data generated by gold standard method. The results of this correlation are visualized by a heatmap (Fig. 4B) and show that the gene expression of each gate (pFACS-RNAseq) on the Y-axis is highly correlated to the expected stage of the gold standard time course (X-axis), demonstrating that pFACS-RNAseq can capture individual stages of *P. falciparum* with high accuracy and reproducibility.

## Discussion

In malaria research, stage specific gene expression profiling throughout the life cycle is essential to better understand parasite biology, drug resistance and phenotypic variation. Unfortunately laboratory culture of *Plasmodium* results in asynchronously staged parasites [27–29] making stage specific studies very challenging. Current methodologies for time series transcriptomics in malaria involve several steps that are time consuming, laborious and may introduce potential confounders such as stress. In this study, we developed a rapid and optimized protocol based on FACS sorting for densely generating stage transcriptome data from asynchronous culture of *P. falciparum* in a more high-throughput manner (Fig. 5). First, we sorted each stage by FACS using live cell dyes, which don’t require fixing cells. We validated the accuracy of this double staining method by sorting each asexual stage and confirming morphological identification by microscopy (Fig. 2C-E). This confirmed that our method can accurately distinguish and sort each developmental stage. The ability to identify and sort clearly discrete stages is a distinct improvement over the time course method using synchronization methods. We then developed an optimized RNAseq protocol for FACS sorted cells. To evaluate the quality of our protocol, we compared the result of library preparations obtained from pFACS-RNAseq to a commercial kit protocol (QIAGEN) and to “gold standard” data obtained from tightly synchronized parasite cultures and library preparation for bulk RNAseq.

**Fig. 5.**
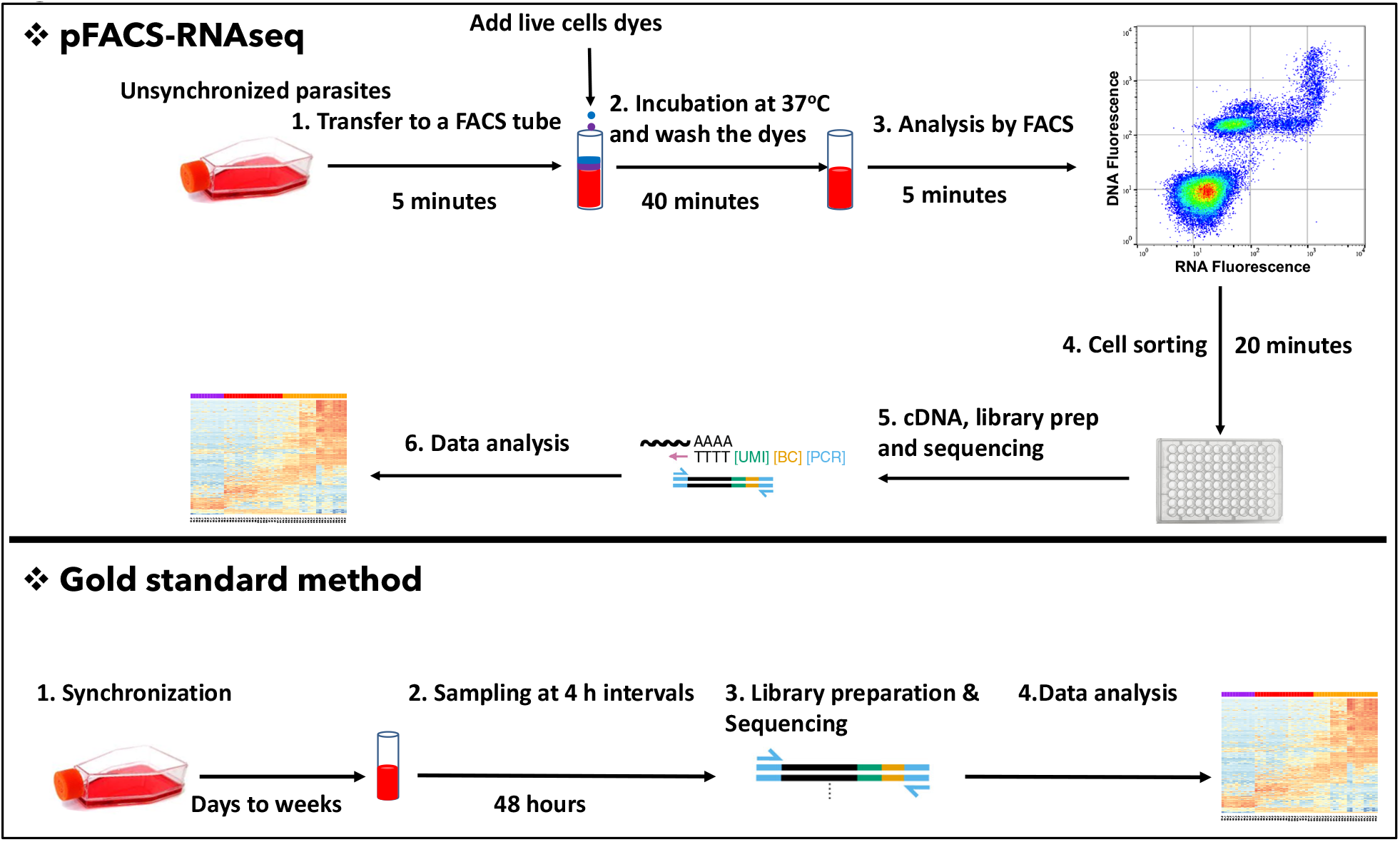
Temporal schematic overview of the pFACS-RNAseq protocol versus gold standard approach for profiling malaria parasites asexual stage specific transcriptomics. For the pFACS-RNAseq, unsynchronized parasites were stained by both DNA and RNA live cell dyes. After 30 minutes of incubation at 37 °C, the dyes are washed with RPMI media and the samples analyzed by FACS. The desired gates were sorted into a 96 well plate for cDNA synthesis. All the sample preparation process before library preparation is done in ~70 minutes. After oligo-dT annealing, template-switching oligo (TSO) is added and 1^st^ and 2^nd^ strand synthesis performed. After this step, all the samples were bead cleaned and PCR amplified for full length cDNA. Library preparation was done from the cDNA with ¼ volume KAPA HyperPlus kits. For the gold standard approach the sample preparation before library preparation can take days to weeks.

The pFACS-RNAseq protocol shows better read quality with a higher percentage of reads mapping to the *P. falciparum* reference genome and more than double the percentage of reads that mapped to a unique locus than the QIAGEN protocol. Compared to our gold standard data, the pFACS-RNAseq has lower percentage of uniquely mapping reads (Table 1, Fig. 2) however the pFACS-RNASeq protocol show negligible rRNA contamination (Fig. S2), lower than the classical approach for the bulk and recently developed protocols for low input transcriptome profiling [8] mainly due to our protocol optimization for a better data quality. This result, show how qualitative is the pFACS-seq because one of the major problems in RNAseq experiments is contamination by rRNA, wasting reagents and lowering the recovery of RNA species of interest [30]. Although pFACS-RNAseq did not outperform the QIAGEN protocol in the number of genes detected, the comparative cost ($15 vs $43), reproducibility and purity of the data are substantially improved.

The malaria parasite IDC has been extensively investigated by transcriptome profiling methods, and these investigations have shown a continuous progression of gene expression changes throughout the IDC. Our global gene expression analysis by MDS, an unsupervised clustering method, shows that we can accurately reproduce the progression of IDC with the pFACS-RNAseq protocol. In fact, 3 mains groups were identified by our clustering method indicating three different transcriptional profiles between these groups. The analysis within groups show that samples belonging to the rings are more similar from one to another and heterogeneity within group increases with the late stages (see axis 1 of the MDS, Fig. 3A, pFACS-RNAseq). These results are in agreement with previous studies suggesting very low transcriptional activity in early stages of the IDC [31]. The profile of gene expression signatures we found (Fig. S3) follows the same pattern that previous stage specific studies described, with gene expression commonly regulated with stages specific genes. Certain genes are highly expressed in early stage, other in middle stage and different group of other genes highly expressed in late stage of the parasite IDC. These observations confirm that the pFACS-RNAseq is an adequate method for malaria stage specific studies.

Gene expression profiling in malaria typically requires large-scale cultures to be generated [32] as well as synchronization and laborious sampling throughout the 48 hours of the IDC. Other methods like 10x single cell RNAseq are increasingly used to explore transcriptomic dynamics throughout different malaria species IDC and could in theory be used to provide stage specific data from unsynchronized parasite cultures. However, these approaches are extremely expensive, and therefore impractical for large scale transcriptomics. More importantly, current 10x transcriptomics protocols produce extremely sparse transcriptomics data from each cell [7, 33]. While these approaches are useful for categorizing parasite to particular lifecycle stages, they provide insufficient resolution for most quantitative genomics applications.

The pFACS-RNAseq method significantly reduces the sampling time from ~6 days to ~2 hours by using only 14μL of asynchronous culture stained with live cell dyes. Although double staining methods were previously used to monitor malaria parasite stage development and growth [17, 18, 34], in our study we fractionated the 3 main stages into multiple gates, then micro-sampled them in a high-throughput manner, and finally confirming with RNAseq profiling that these sorted gates correspond to tightly synchronized samples of IDC progression.

The fact that the gold standard method detects higher number of genes than the FACS sorted methods might have a number of explanations. Firstly, in our study, for the gold standard approach, each sample was generated from millions of cells while the FACS-sorted cells were collected from 200-3,000 cells. Secondly, in our study, the gold standard samples were sequenced deeper that our FACS-sorted samples. Crucially, the gold standard samples were generated using standard synchronization approaches, we cannot discount the presence of all intraerythrocytic life cycle stages at each time point at a low level artificially increasing the breadth of genes detected.

pFACS-RNAseq has many advantages compared to the gold standard approach. pFACS-RNAseq demands less culture work, doesn’t require multiple rounds of synchronization of parasite cultures and eliminates the laborious 48 hours of sampling. The general protocol is highly flexible. For instance, it can be limited to specific life cycle stages, the base RNA amplification approach can be altered as novel protocols become available and the numbers of cells can be both increased and decreased to provide either improved detection of heterogeneity for single cells, or improved coverage and quality for larger cell populations. We believe that these features of pFACS-RNAseq will make it broadly applicable for high throughput stage-specific studies of *P. falciparum.* Furthermore, this approach will also make transcriptomic analysis of other *Plasmodium* species like *P. vivax,* for which long term culture is not possible, and will also be applicable for many other organisms to explore transcriptional dynamics from asynchronous cultures.

## Materials and methods

### Cell culture, staining and flow cytometry sorting

Unsynchronized cultures were grown to 7-9% parasitemia in standard conditions (CM:RPMI 1640, with L-glutamine, 25mM HEPES, gentamicin, 0.5% AlbuMAXII, and 50ug/mL hypoxanthine with type O+ donor RBCs at 4% hematocrit), and harvested by centrifugation (1600 rpm/4 minutes). Fourteen μl of the cell pellet were added to a tube containing 4 mL of 1X RPMI/HEPES/gentamicin solution without AlbumaxII/hypoxanthine (ICM) with 1.5 μl of VybrantDye Violet™and 1.5 μl of SYTO RNA Select™Green (both from ThermoFisher). This solution was incubated in the dark at 37° C, shaking, for 30 min. After incubation, cells were washed twice in warm ICM and resuspended in 6 ml of ICM prior to flow cytometry. The samples were analyzed on a BD Influx cytometer (BD Biosciences, San Jose, Ca, USA) equipped with a 100μm nozzle. Cells were gated based on their DNA and RNA fluorescence and 50,000 events were acquired. The SYTO RNA™stain was excited with a blue laser (488 nm) and the 530/40 band pass filter was used to collect the emitted light. The DNA fluorescence was detected by the 355 nm UV laser with the 460/50 band pass filter. The gating strategy was carried out from tests on stained samples prepared from infected and uninfected red blood cells (data not shown).

Each stage gate (ring, trophozoite, schizont) was sorted and thin smears were stained with Wright-Giesma for examination by microscopy in order to confirm that morphology of the sorted gate corresponded to the expected stage.

### Gold standard time course

Parasites were grown to over 5% parasitaemia in 100mL CM with 4% hematocrit. Cultures were tightly synchronized using 3 sorbitol treatments. The first synchronization occurred when the parasites were mostly rings, then a second synchronization was performed 10 hours later. A final sorbitol synchronization was performed when parasites reinvaded the red blood cells and were early rings, 18 hours after the second synchronization. To minimize stress on the parasites during RNA collection, parasites were placed into 6-well plates, each containing 5ml of culture, at 2% hematocrit. Each sample and technical replicate was collected every 4 hours, for 56 hours total. The parasitemia prior to reinvasion was 2% for ring stage time points, and 1% parasitemia for trophozoite and schizont time points. When parasites began to reinvade the red blood cells and were 90% early rings, collections for RNA began as Time 0. Each well was collected separately and washed with 1x PBS, then 1ml of TRIzol Reagent (ThermoFisher) was added to each pelleted sample and homogenized with needle and syringe. Samples were mixed at 37°C for 5 minutes, then stored at −80°C. RNA was extracted using Direct-Zol RNA Mini Prep (Zymo Research), quantified with Qubit RNA BR Assay Kit (ThermoFisher), and quality assessed with Bioanalyzer RNA 6000 Nano assay (Agilent). RNA libraries were prepared, according to manufacturer’s directions, with 500ng of total RNA, using the KAPA Stranded mRNA-Seq Kit (KAPA Biosystems), with 7 PCR cycles. Samples were pooled by nM equivalents from each reaction and sequenced on an Illumina HiSeq 2500 with 2×100 flow cells.

### QIAGEN FX Single Cell RNA Library Kit (QIAGEN)

used according to manufacturer’s directions. Briefly, 100 cells per gate were collected in 96 well Lo-Bind plates (Eppendorf, Hauppage, NY) containing 5uL of 1XPBS (Lonza) per well using stringent precautions against contamination [22]. Samples were placed on dry ice, and then stored at −80°C until cDNA creation. 50ug of whole transcriptome amplified (WTA) cDNA were fragmented and adapter ligated according to directions except a 20-minute fragmentation incubation at 32°C was used. Ligated cDNA was measured by QBit Broad Range DNA Kit (Invitrogen) using 2uL of the product, and 1uL was run on the Agilent 4200 Tape Station (Agilent) for estimating fragment size and product purity. To improve yield and resolve correct fragment size, samples were amplified by 8 cycles of PCR with the QIAGEN Gene Read DNA Amp Kit.

### pFACS-RNAseq Development Protocol

Adapted from the molecular crowding single-cell RNA sequencing (mcSCRB-seq) protocol [25]. Cells were sorted into 5uL lysis buffer containing 0.8% Triton-X100 and 2U SUPERase Inhibitor (Invitrogen). During the development phase, various cell numbers were tested from 3000 to single cell inputs. Final cell numbers were 3000 for each gate in the ring stage, 200 cells for each gate in the trophozoite stage, and 200 for each gate in the schizont stage. For samples from ring gates we added 3.25uL of 10uM oligo-dT/dNTP per well due to increased sort volume for high cell numbers; for trophozoite and schizont gates we added 1.0uL oligo-dT ACACTCTTTCCCTACACGACGCTCTTC **CATATTCCTGGTGG**NNNNNNNN T_24_ (Eurofins) after thawing and prior to denaturation. The sequence in bold font indicates the 14bp barcode used in the Quartz-Seq 2 protocol [26]. The reverse transcription (RT) Master Mix contained 2.2uL of 5X Maximus H-Reaction Buffer (ThermoFisher), 1.65uL of 50% PEG8000 (Fisher), 0.44uL of 25mM dNTP (ThermoFisher), 0.11uL of the 100uM TSO primer ACACTCTTTCCCTACACGACGC (G)(G)(G) (Eurofins), and 0.11uL of Maximus H-RT (ThermoFisher). For samples from ring gates we added 17.9uL of RT Master Mix (in the same proportions) to the denatured RNA/Oligo-dT sample while trophozoite and schizont samples received 5.5uL. Samples were centrifuged, tip-mixed gently but thoroughly, and centrifuged again. cDNA synthesis was carried out at 42°C for 90 min. Samples were then centrifuged, bead cleaned with 1X AMPure beads (30%PEG) and eluted in 12uL of EB. Finally, samples (10uL) were amplified using 15uL of PCR Master Mix containing 12.5uL 2X Terra Buffer (ThermoFisher), 0.5ul 10uM IS-PCR primer ACACTCTTTCCCTACACGACGC (Eurofins), 0.5uL Terra Polymerase (ThermoFisher), and 1.5uL ultra-pure water. The final PCR program used was: 1 cycle of 98°C for 3 min, 19-36 cycles of (98°C/15 sec, 65°C/30 sec, 68°C/4 min), followed by 1 cycle of 72°C for 10 minutes. The PCR reaction was bead cleaned using 0.8X AMPure beads and eluted in 18uL EB. Samples were measured for cDNA using the QBit BR DNA kit. Barcode/UMI sequences were adapted from the Quartz-Seq2 protocol [26], and the oligo-dT length was shortened for the mcSCRB protocol for use with *Plasmodium falciparum.*

### qPCR Protocol

We used qPCR to determine the number amplification cycles needed for different number cells for each sorted gate from the 3 mains stage of P. falciparum. We followed the pFACS-RNAseq Development Protocol above except for the PCR amplification phase. Here we performed ½ reaction volumes and added 0.5uL of a 1/1000 dilution of SYBR Green (ThermoFisher) and only 0.25uL water. We ran the QPCR reaction on the QuantStudio 5 (Applied Biosystems) with the same PCR protocol.

### KAPA Library Preparation and Sequencing

pFACS cDNA libraries were prepared with the KAPA HyperPlus Kit according to manufacturer’s directions using 50ng of cDNA and ¼ volume reactions. Small modifications were made including ligation for 1 hour, amplifying for 7 cycles, using size selection after amplification, and bringing up the volume of sample to full volume before the first cut of the size selection. We used the KAPA Dual-Indexed Adapter Kit, adding 7.5uM adapter to the appropriate well. Individual samples were measured for DNA quantity using the QBit BR DNA Kit. Samples were then pooled for sequencing based on their QBit measurements to normalize input. The pooled sample was quantified using the KAPA Library Quantification Kit and adjusted to 2-4nM with Buffer EB (QIAGEN) for sequencing on Illumina platforms. The pool was also run on the Agilent Tape Station using the D1000 Kit to assess fragment size and quality. Pools were run on the Illumina HiSeq 2500 or Illumina NextSeq for 2×150bp runs.

### RNA-seq analysis

All FASTQ files were trimmed using fastp [35] to reduce impact of lower quality reads and the read quality was checked using the FastQC program. Trimmed sequences were aligned to the PF3D7 reference genome using STAR-2.5.3 [36]. SAM files were converted to BAM using samtools v-1-3 view −b and sorted with samtools v-1.3-n. The total gene count was determined using HTSeq v-0.7.2 [37] with htseq count. Read count per million (CPM) were calculated for the downstream analysis. For the differential gene expression (DGE) comparison, we first matched up the QIAGEN and pFACS-RNAseq data on the gold standard time course. Then we selected tight windows (3 hours intervals) for the analysis. All the data for DGE analysis is filtered by mean reads depth threshold of 10 across the retained data. In order to identify co-expressed gene clusters, a weighted gene co-expression network analysis (WGCNA) was constructed using gene expression data from 13 time points (gold standard data). All genes (5360) we detected by this standard approach were usded the gene co-expression network construction, with the parameters of networkType = signed, softPower = 13 and minModuleSize = 100. The gene expression level (normalized by gene) is plotted using a heatmap All the plots from the downstream analysis were generated by R (v-3.6.3).

### Statistical analysis

Performances of the approaches (gold standard, pFACS-RNAseq and QIAGEN) in our study were first evaluated by mapping statistics. We performed Kruskal-Wallis tests to examine if the observed differences in uniquely mapping reads, in multimapping reads and in unmapped reads between approaches are significant. As the Kruskal-Wallis tests will indicate a global comparison, we also computed a pairwise Wilcox test for multiple comparison between methods with Benjamini-Hochberg adjusted *P-value* [38].

## Supporting information

Combined Supplementary Figures, Text and Tables

## Acknowledgments

We thank member of the Ferdig, Vaughan, Anderson and Cheeseman laboratories for assistance during this project. This work was funded by National Institutes for Health (https://www.nih.gov) grant P01 AI127338 (to Michael Ferdig), NIH grants NIAID R01 AI110941-01A1 to IHC and NIH grant R37 AI048071 to TJCA. IHC is a Milton S. and Geraldine M. Goldstein Young Scientist.

