## Supplementary material for "Efficient transcriptome profiling across the malaria parasite erythrocytic cycle by flow sorting": Combined Supplementary Figures, Text and Tables

### Supporting information

**Fig. S1***. Boxplot representing linear correlations between replicates from the QIAGEN protocol. The correlations between replicates were estimated by Pearson method.*

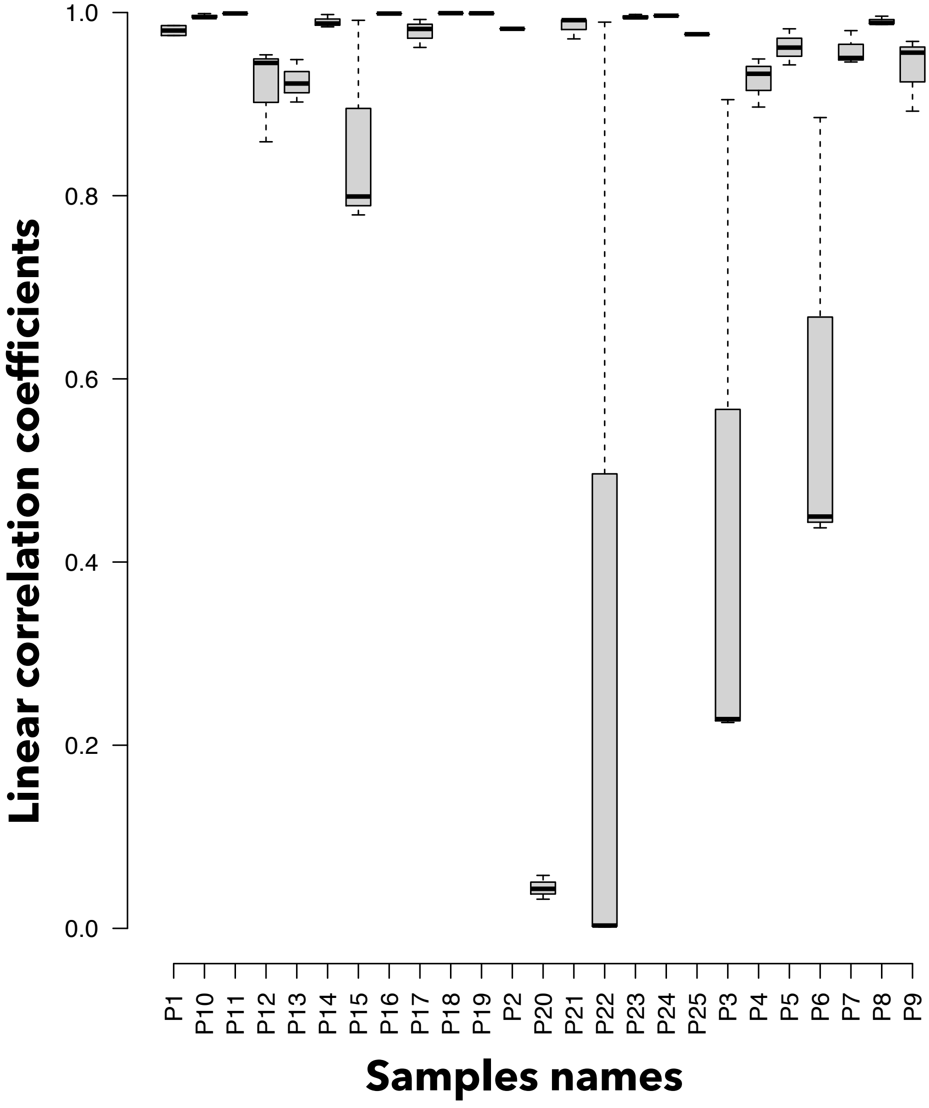

***Fig. S2.*** *Summary of read counts* ***showing the*** *proportion of reads mapping to P. falciparum coding genes, no feature, ribosomal RNA; multi-mapping read and unmapped reads for the gold-standard, the Qiagen and the pFACS-RNAseq protocols.*

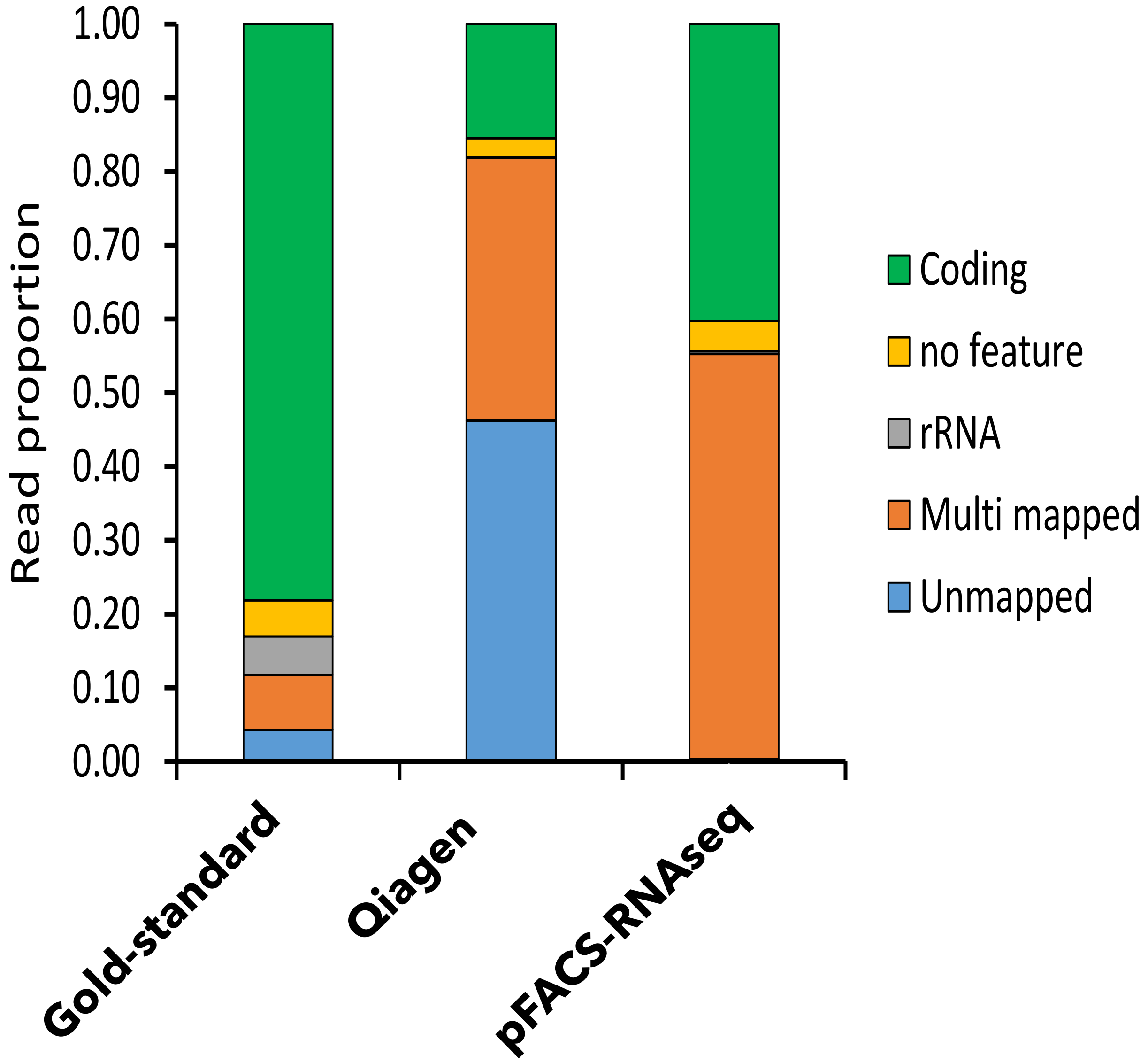

**Fig. S3.** *The top 350 most expressed genes revealing different patterns of gene expression throughout the intraerythrocytic development cycle (dark blue: low gene expression and dark orange: high gene expression).*

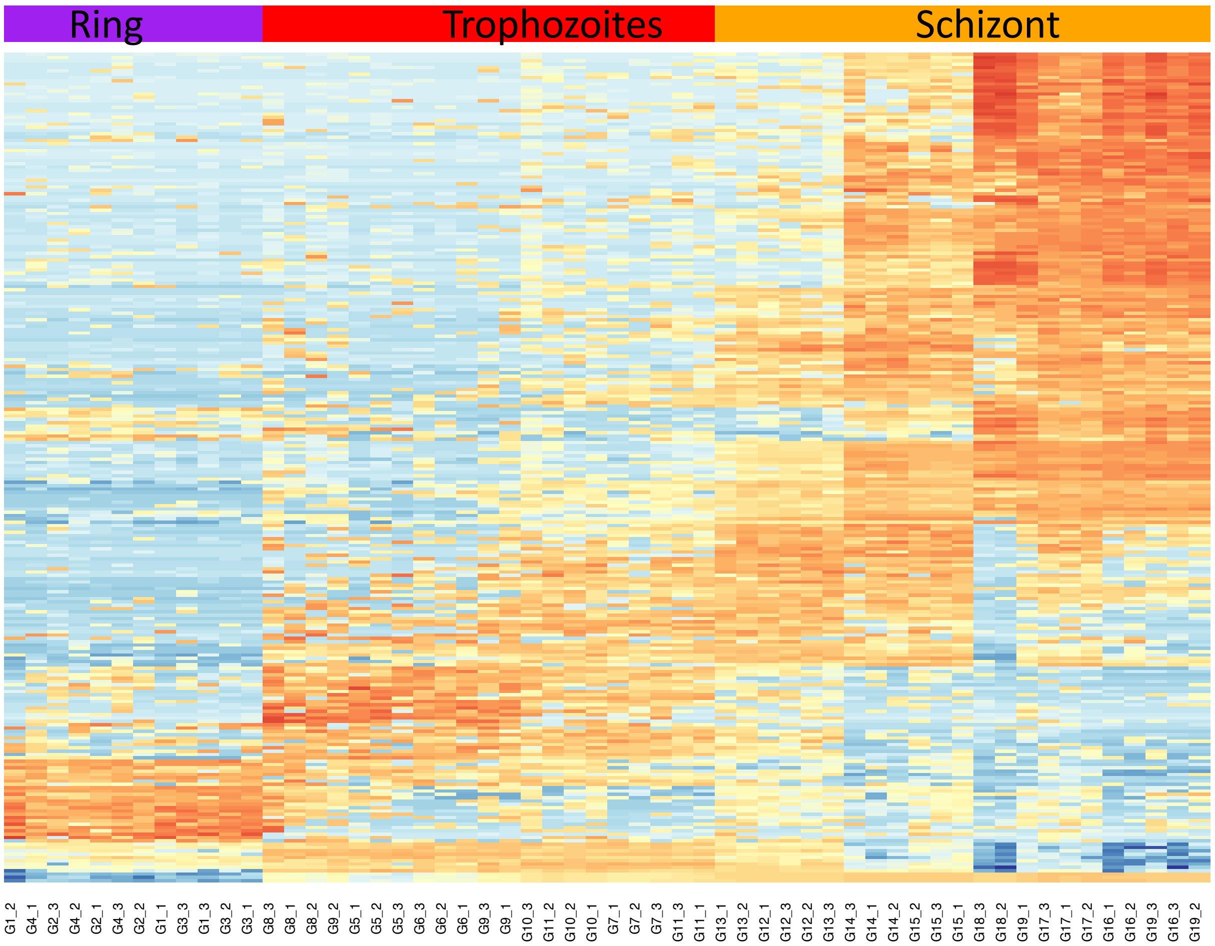

**Fig. S4**. QPCR results of differing parasite number per reaction

A. Rings

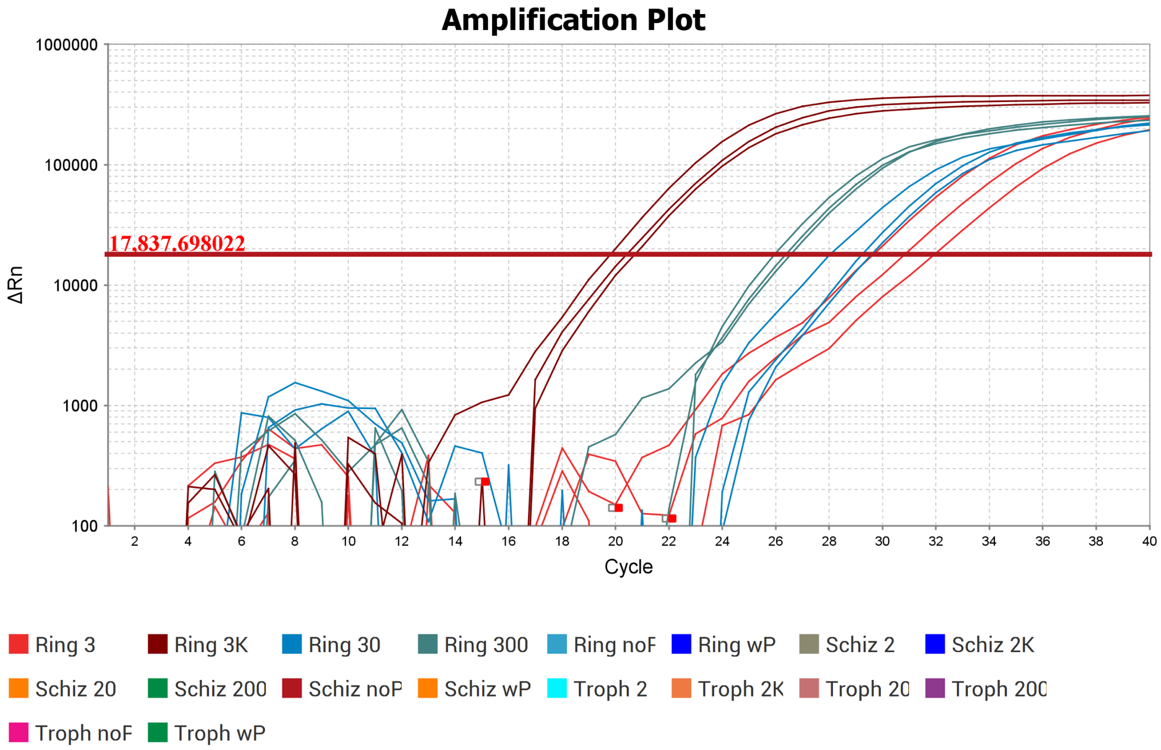


B. Trophozoites

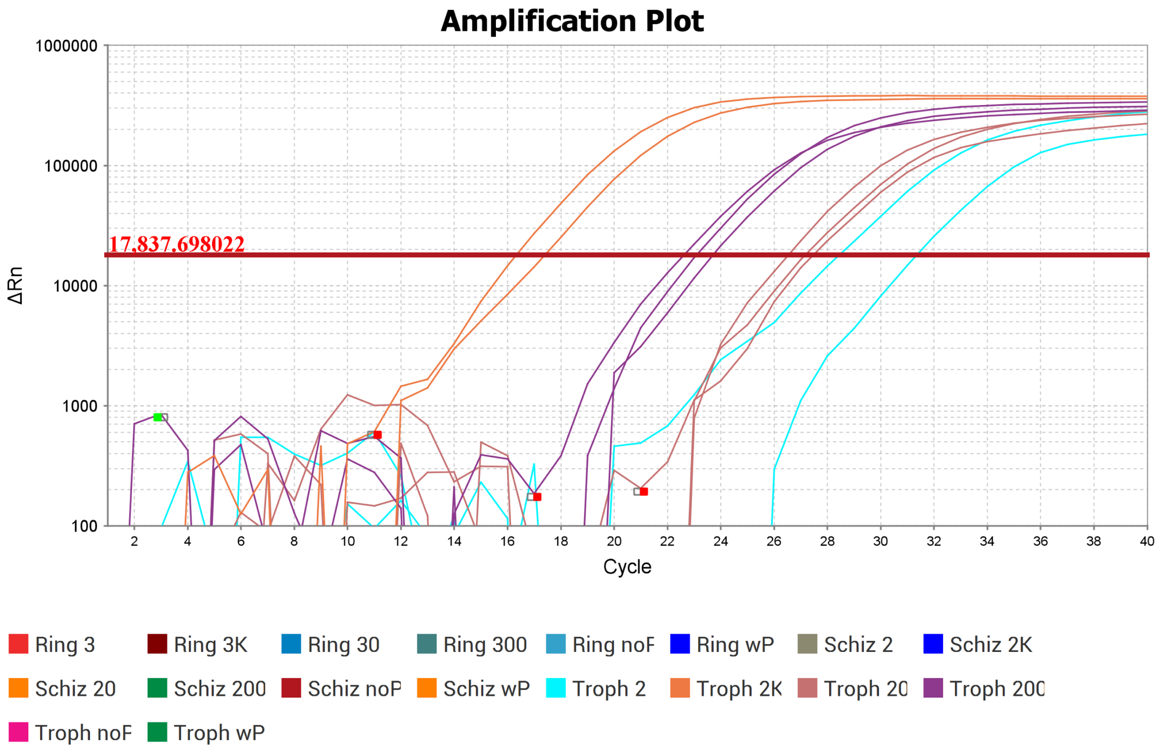

C. Schizonts

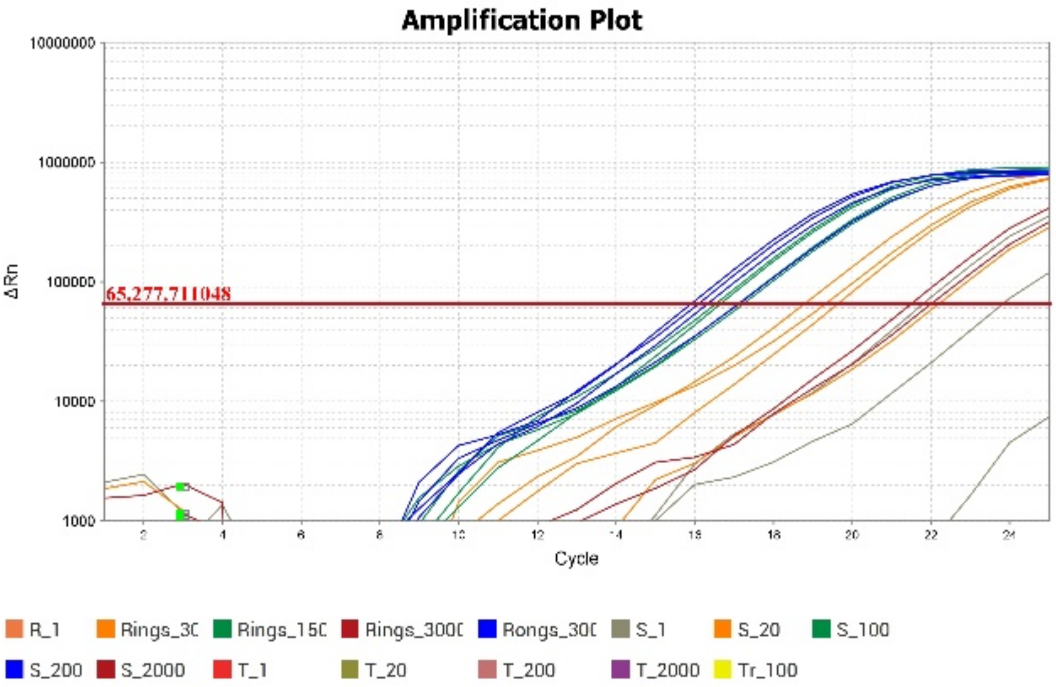
Amplification Plot
10000000
1000000
100000
10000
1000
ΔRn
65,277.711048
Cycle
R_1
Rings_3C
Rings_15C
Rings_300C
Rongs_30C
S_1
S_20
S_100
S_200
S_2000
T_1
T_20
T_200
T_2000
Tr_100

**Table S1:** *The Kruskal-Wallis test between uniquely mapping reads for the 3 methods is significant (Kruskal-Wallis chi-squared = 127.05, df = 2, p-value < 2.2e-16). We computed a pairwise Wilcox test for multiple comparison between methods with Benjamini-Hochberg adjusted P value [1] for the uniquely mapping reads.*

| | pFACS-RNAseq | Gold standard |
|---|---|---|
| Gold standard | 4.2e-13 | NA |
| QIAGEN | < 2e-16 | 6.2e-14 |

**Table S2.** *The Kruskal-Wallis test between multi-mapped reads for the 3 methods is significant (Kruskal-Wallis chi-squared = 120.02, df = 2, p-value < 2.2e-16). We computed a pairwise Wilcox test for multiple comparison between methods with Benjamini-Hochberg adjusted P value [1] for the uniquely mapping reads.*

| | pFACS-RNAseq | Gold standard |
|---|---|---|
| Gold standard | 6.3e-13 | NA |
| QIAGEN | < 2e-16 | 9.2e-11 |

**Table S3.** *The Kruskal-Wallis test between unmapped reads for the 3 methods is significant (Kruskal-Wallis chi-squared = 129.55, df = 2, p-value < 2.2e-16). We computed a pairwise Wilcox test for multiple comparison between methods with Benjamini-Hochberg adjusted P value [1] for the uniquely mapping reads*

| | pFACS-RNAseq | Gold standard |
|---|---|---|
| Gold standard | 2.1e-12 | NA |
| QIAGEN | < 2e-16 | 7.5e-14 |

**Table S4. Kits, methods, and reagents used during optimization of mcFACS-Seq.** **oligo-dT sequence adapted from Sasagawa et al. 2018 [2].*

| Condition | QIAGEN FX | Reid [3] (SmartSeq) | Bagnoli [4] (mcSCRB) | Test 1 Reid | Test 2 (pFACS-RNAseq) | Test 3 (pFACS-RNAseq) |
|---|---|---|---|---|---|---|
| **Lysis Buff in sort PL** | 1X PBS | 0.8% Triton X-100 | PHusion, PK | 0.8% Triton X-100 | 0.8% Triton X-100 | 0.8% Triton X-100 |
| **Oligo-dT** | Random hexamer | 25–T, non-anchored | 30-T, anchored | 25–T, anchored | 25–T, non-anchored | 25–T, non-anchored QS2 |
| **RT additives** | NA | $MgCl_2$, betaine | none | $MgCl_2$, betaine | none | none |
| **RT enzyme** | QIAGEN | SmartScribe | Maximus H- | SmartScribe | Maximus H- | Maximus H- |
| **PCR enzyme** | QIAGEN | KAPA HotStart | Terra | KAPA HotStart | Terra | Terra |
| **PCR Cycles** | WTA | 25 | 15 | 25 | 25 | 30 |

***Table S5.*** *Effect of adding PEG to the reverse transcription step*

| Cell Number | Stage | QPCR $C_T$ without PEG | QPCR $C_T$ with PEG |
|---|---|---|---|
| 3000 | Ring | 12.3 | 10.53 |
| 200 | Trophozoite | 10.7 | 8.34 |
| 200 | Schizonts | 15.6 | 10.5 |

### Supplementary Text

#### Early method development

We tried several iterations of published protocols [3, 4] and largely used quantification by Qbit and/or QPCR to determine best practices for our final method listed in this paper (pFACS-RNAseq). We began with the Reid et al 2018 [3] paper and after some changes for our laboratory to use existing library production, we proceeded to sequence a few samples.

**Reid Adapted Protocol**: Following Reid's protocol [3] we collected 100 cells per gate into 5uL of 0.8% Triton-X100 containing 2U SUPERase Inhibitor (Invitrogen). Samples were placed on dry ice and then stored at -80°C until cDNA creation. Plates were thawed briefly on ice, centrifuged for 30 sec/750xG, and 2.5uM oligo-dT AGCAGTGGTATCAACGCAGAGTACT$_{30}$VN (Eurofins) along with 2.5mM dNTP (ThermoFisher) were added. The samples were denatured at 72°C for 3 minutes, followed by snap cooling on ice. RT Master Mix was prepared containing 1X RT reaction buffer (TaKaRa), 0.5U SUPERase Inhibitor (Invitrogen), 1 uM TSO primer AGCAGTGGTATCAACGCAGAGTACATrGrG+G (QIAGEN), 6uM $MgCl_2$ (ThermoFisher), 1M betaine (Affymetrix), 50uM DTT (TaKaRa), 0.5uL of SMARTScribe RT (TaKaRa) and ultra-pure water (QIAGEN) for volume – 5.5uL of RT Master Mix was added to each well of the denatured sample created above. First strand cDNA was synthesized by incubation with the following program: 1 cycle of 42$^0$C for 90 minutes, 10 cycles of (42°C/2 min, 50°C/2 min), and a final cycle of 70°C for 15 minutes. Samples were removed and snap cooled on ice, then centrifuged at 750xG/30 seconds. Finally, samples were amplified by adding 15uL of PCR Master Mix: 1X KAPA HiFi Ready Mix (KAPA Biosystems/Roche), 2.5 uM IS-PCR primer AAGCAGTGGTATCAACGCAGAGT (QIAGEN), and ultra-pure water to volume. Samples were amplified with the following program: 1 cycle of 98°C for 3 min, 25 cycles of (98°C/20 sec, 67°C/15 sec, 72°C/6 min), followed by 1 cycle of 72°C for 5 minutes. After amplification, samples were centrifuged as above, and bead cleaned with 1X AMPure Beads (Beckman Coulter) and eluted in 14ul of Buffer EB (QIAGEN). Samples were measured for cDNA using the QBit BR DNA kit.

We modified this protocol for qualitative improvement by trying the mcSCRB-Seq [4] (see materials and method for the pFACS-RNAseq development protocol).

**Early development results**
Supplementary table 5 provides an overview of the original methods and our method development conditions. Early sequencing results from the Reid et al. 2018 protocol included a large amount of rRNA reads (data not shown) so we went forward with developing the mcSCRB protocol. The differences between Test 1 [3] from Reid et al. 2018 and Test 2 (pFACS-RNAseq) included the change to the reverse transcription master mix of Bagnoli et al. 2018 [4] and using the Bagnoli BC-UMI oligo-dT. The result of these full-length oligos did not seem to work. Thus Test 3 was performed with shortened oligo-dTs based on sequences from the Sasagawa et al. 2018 paper [2] to improve yield and decrease bias.

We investigated whether the addition of PEG to the RT reaction would improve our yield, we used the Reid oligo-dT for this test. We observed a definite increase in yield as evidenced by the lowering of the $C_T$ value in a QPCR reaction. Representative curves of the QPCR reaction can be seen in Fig. S4.
